# Distinct evolutionary paths in chronic lymphocytic leukemia during resistance to graft-versus-leukemia

**DOI:** 10.1101/2020.04.09.033555

**Authors:** Pavan Bachireddy, Christina Ennis, Vinhkhang N. Nguyen, Kendell Clement, Satyen H. Gohil, Sachet A. Shukla, Juliet Forman, Nikolas Barkas, Samuel Freeman, Natalie Bavli, Liudmila Elagina, Ignaty Leshchiner, Arman W. Mohammad, Laura Z Rassenti, Thomas J Kipps, Jennifer R. Brown, Gad A. Getz, Vincent T. Ho, Andreas Gnirke, Donna Neuberg, Robert J. Soiffer, Jerome Ritz, Edwin P. Alyea, Peter V. Kharchenko, Catherine J. Wu

## Abstract

Resistance to the graft-versus-leukemia (GvL) effect remains the major barrier to successful allogeneic hematopoietic stem cell transplantation (allo-HSCT) for aggressive hematologic malignancies. The basis of GvL resistance for advanced lymphoid malignancies remains incompletely understood. We hypothesized that for patients with chronic lymphocytic leukemia (CLL) treated with allo-HSCT, leukemic cell-intrinsic features shape GvL outcomes by directing the evolutionary trajectories of CLL cells. Integrated genetic, transcriptomic and epigenetic analyses of CLL cells from 10 patients revealed that the clinical kinetics of post-HSCT relapse are shaped by distinct molecular dynamics and suggest that the selection pressures of the GvL bottleneck are unlike those imposed by chemotherapy. No selective advantage for HLA loss was observed, even when present in pre-transplant subpopulations. Regardless of post-transplant relapse kinetics, gain of stem cell modules was a common signature associated with leukemia relapse. These data elucidate the biological pathways that underlie GvL resistance and post-transplant relapse.

**One Sentence Summary:** We find that the clinical kinetics of chronic lymphocytic leukemia relapse after stem cell transplant are underwritten by distinct genetic and epigenetic evolutionary trajectories and suggest that the selection pressures of the post-transplant, immunologic bottleneck are unlike those imposed by chemotherapy.

## Introduction

Allogeneic hematopoietic stem cell transplantation (allo-HSCT) is one of the earliest forms of successful cancer immunotherapy whose study has elucidated critical insights into tumor-immune interactions (*1, 2*). As the only curative option for aggressive and advanced hematologic malignancies, allo-HSCT derives its potency from the underlying, donor-derived graft-versus-leukemia (GvL) effect wherein donor immune cells recognize and eradicate recipient leukemic cells. Reduced-intensity conditioning regimens (RIC) for allo-HSCT are increasingly used to reduce treatment-related morbidity and preserve the curative GvL effect (*3, 4*), but post-transplant disease recurrence limits their efficacy. Therefore, elucidating the mechanisms underpinning GvL resistance is vital to improve transplant outcomes.

Multiple studies have identified various pathways of GvL resistance associated with relapse of myeloid malignancies, including loss of HLA class I and II genes, upregulation of inhibitory immune checkpoint molecules, and oncogene-driven immune evasion (*5–9*). For lymphoid malignancies however, less is known about potential mechanisms of GvL evasion. In particular, how underlying genetic and epigenetic features inform the clinical kinetics of post-transplant relapse or whether dysregulation of HLA class I and II genes influences GvL resistance remains unclear.

To address these questions, we assembled a cohort of 10 patients with chronic lymphocytic leukemia (CLL), the majority of whom were treated with RIC allo-HSCT. We hypothesized that leukemic-intrinsic features primarily shape the evolutionary dynamics of CLL cells during GvL resistance. We therefore focused on the evaluation of longitudinal changes in genomic, epigenomic and bulk and single cell transcriptomic features of CLL cells following transplant. By defining the genetic, epigenetic and transcriptomic changes that shape CLL relapse after allo-HSCT, we demonstrate that these evolutionary paths are unique to the GvL immunologic bottleneck and identify the underlying mechanisms that influence the clinical kinetics of post-HSCT relapse.

## Results

### Clinical kinetics of CLL relapse after allo-HSCT correspond to distinct evolutionary paths

We assembled a cohort of 10 CLL patients with varying time to progression after allo-HSCT (range: 83-1825 days), for whom paired pre- and post-transplant relapse specimens were available (see **Methods**). As expected for patients with CLL undergoing allo-HSCT, patients had received multiple lines of therapy prior to transplant (median: 3, range: 1-6) and 6 of 9 assessable patients had unmutated IGHV status (**Figure 1A, Table S1**).

**Fig. 1.**
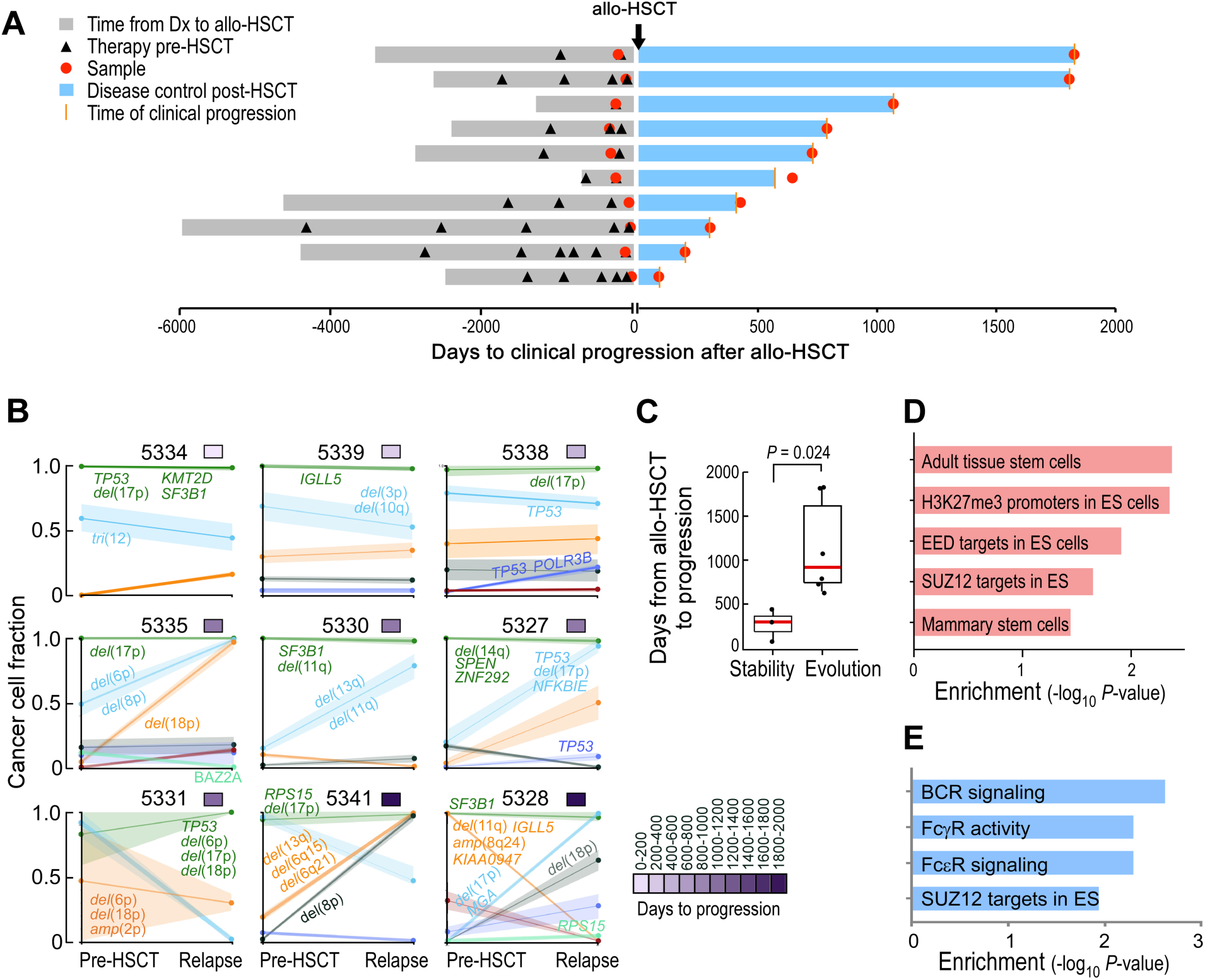
Timing of relapse after allo-HSCT is defined by distinct evolutionary trajectories and stem cell expression programs. (**A**) Time to clinical progression after allo-HSCT for 10 patients with CLL indicated along with relevant clinical histories and times of sample procurement. (**B**) Evolutionary patterns of mutation clusters shown by their cancer cell fractions (CCFs), inferred using PhylogicNDT. Time to progression is shown by the purple color bar. Putative CLL drivers detected for each cluster are shown. (**C**) Box plot of time to relapse after allo-HSCT in patients with (n=6) and without (n=3) clonal evolution. The *P* value was calculated by a two-sided Wilcoxon ranked sum test. “Evolution” was defined as having any cluster with absolute difference ≥0.2 between pre-HSCT and relapse timepoints (see **Methods**). (**D**) Unadjusted *P* values of enriched stem cell gene sets in pretreatment allo-HSCT samples per GSEA, comparing samples collected from early versus late relapses. (**E**) Unadjusted *P* values of enriched signaling pathways per GSEA, comparing post- versus pre-HSCT samples of late relapses.

We performed whole-exome sequencing (WES) of DNA isolated from purified CLL cells from paired pre- and post-transplant relapse samples (see **Methods**) and matched donor and recipient germline DNA from 9 of 10 patients (median coverage of 160x; **Table S2**). Compared to treatment naïve CLL (*10*), we detected a greater number of non-silent single nucleotide variants (sSNVs) and somatic insertions and deletions (sIndels) per pre- and post-transplant exome (a median of 39 and 41), respectively (p<0.01; **Figure S1A, Table S3, Methods**), consistent with the extensive chemotherapeutic exposure in patients undergoing allo-HSCT. Nevertheless, the spectrum of CLL cancer drivers in sSNVs, sIndels, and copy number alterations (CNAs) was similar to that previously observed (*10*) (**Figure S1B, Table S4**). Consistent with the aggressive nature of these leukemias, we observed multiple patients to have mutations and CNAs involving *TP53* and *SF3B1*, but found no somatic alterations uniquely shared among post-transplant samples.

To examine patterns of clonal dynamics associated with relapse, we used the tool PhylogicNDT (*11, 12*), which reconstructs phylogenetic and evolutionary trajectories of cells based on integration of somatic alterations detected from multiple samples. We clustered mutations together based on their cancer cell fractions (CCFs) and examined their change over time. Although all patients displayed multiple and diverse CLL drivers at baseline (**Figure S1C, Table S5**), their evolutionary dynamics during relapse segregated into two distinct patterns: clonal stability (defined by changes in all cluster CCFs <0.2) and clonal evolution (changes in any cluster CCF ≥0.2) (**Figure 1B**). We found that those samples with clonal stability originated from the 3 patients with the shortest times to relapse (median of 304 days; ‘early’ relapse), while clonal evolution was evident in the samples from the 6 patients with longer times to relapse (median of 798 days; ‘late’ relapse, p=0.024**) (Figure 1C**). We also found *in silico* evidence for GvL immune activity in the late relapsing patients. Predicted neoantigens with strong binding affinity (**Methods**; defined as binding affinity ≤50nM (*13, 14*)) were enriched in CLL clonal clusters that contracted post-transplant (present in 25% of contracting [see **Methods**], 4% in stable and 7% in expanding mutations; p<0.001, **Figures S1D-S1E, Table S6**). Additionally, contracting neoantigens overall had stronger predicted binding affinities than those that either expanded or remained unchanged (p=0.04; p=0.00014, respectively; **Figure S1F**) and were observed only in late relapsers, including patients 5328, 5331, and 5335 (**Figure S1G**).

To investigate the pre-transplant molecular profiles that may herald clonal stability or evolution, we performed differential gene expression analysis between pre-transplant CLL cells from early versus late relapsers and found 853 upregulated and 793 downregulated genes (FDR<0.25). Gene set enrichment analysis (GSEA) highlighted stem cell modules in the early relapses, suggesting the presence of stem cell-like states in the pre-transplant setting to be important for subsequent rapid relapse, particularly components of the polycomb-repressive complex 2 (PRC2) such as the *EED* and *SUZ12* gene pathways (*15*) (**Figure 1F, Table S7**). Consistent with the lack of genetic evolution during early relapse, we found only 66 differentially expressed genes between pre- and post-HSCT samples in early relapses suggesting little transcriptional change. However, paired differential expression analysis between pre- and post-HSCT samples in late relapses revealed 1002 differentially expressed genes (FDR<0.25) and upregulation of similar stem cell pathways in addition to Fc and B cell receptor (BCR) signaling (**Figure 1G, Table S8**). Altogether, these data support the notion that early CLL relapse after transplant is characterized by a pre-existing transcriptional state conferring resistance and comprising stem cell properties. This state therefore does not require evolution of the clonal architecture, and subsequent relapse manifests as genetic stability. In contrast, late CLL relapse, occurring after immune reconstitution, is likely subjected to a GvL selection pressure, manifested by neoantigen depletion. This immunologic bottleneck leads to acquired resistance, genetically, via clonal replacement and, transcriptionally, via upregulation of stem cell and FcR/BCR signaling pathways.

### Genetic evolution of CLL cells carries phenotypic consequences

To evaluate the functional consequences of this genetic evolution, we sought to measure changes in associated gene expression of these heterogeneous clonal populations. We adapted a droplet microfluidic-based platform (“inDrops”, **Methods**)(*16*) to obtain single cell transcriptome (scRNA-seq) data from PBMC samples collected from pre- and post-HSCT samples from 2 patients with early relapse and clonal stability (patients 5339 and 5338, relapsing 304 and 443 days after allo-HSCT) and 2 patients with late relapse and clonal evolution (patients 5341 and 5328, relapsing 1801 and 1825 days after allo-HSCT) (**Figure 2A, Table S9**). We profiled a median of 1,261 CLL cells (range: 1,035-3,751) per sample as assessed by *CD19* and *CD5* gene co-expression. The samples were integrated computationally using Conos (*17*) to overcome inter-patient variability and identify major cell populations (**Figure 2B, Figure S2B**).

**Fig. 2.**
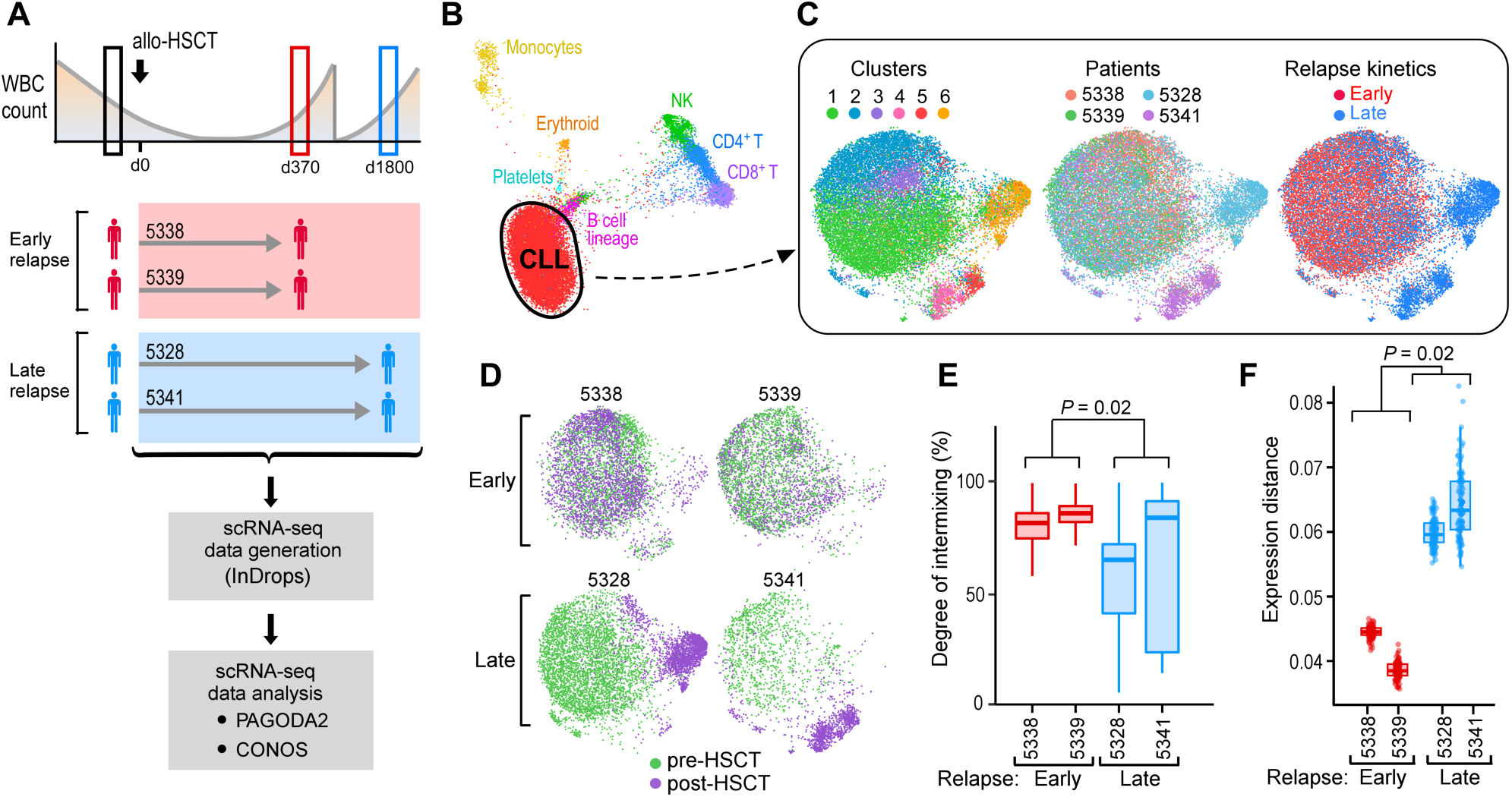
Phenotypic changes in relation to the kinetics of genetic evolution. (**A**) Sample schematic and experimental workflow for capturing single CLL cell transcriptomes through the inDrops system from paired, pre- and post-HSCT PBMCs from two early and two late relapses. (**B**) Joint graph visualized using largeVis embedding showing clustering and annotation of cell subsets from PBMCs from all four patients. (**C**) Joint graphs using UMAP embedding of computationally identified CD19^+^CD5^+^ CLL cells, colored by cluster (left), patient (center) and relapse kinetics (right). (**D**) Individual joint graphs for both early and late relapse patients colored by timing. (**E**) kNN-based quantification of timepoint intermixing across clusters for each patient. The *P* value was calculated from a two-sided Welch t-test comparing means of individual cell values per patient, grouped into early (n=2) versus late (n=2) relapses. (**F**) Extent of gene expression change between pre- and post-transplant CLL cells for each patient. The *P* value was calculated from a one-sided Welch t-test.

The CLL cells comprised 6 transcriptionally distinct clusters that segregated by patient and timing (**Figure 2C**). For the patients who relapsed early, pre- and post-transplant CLL cells exhibited an indistinct population structure, evident by a high degree of intermixing of cells within clusters and consistent with their genetic and transcriptional stability seen on the bulk sample level (**Figure 2D**). In contrast, analysis of late relapsers revealed marked spatial segregation by timepoint with a defined population substructure evident in distinct post-transplant clusters from both late relapses. Late relapse CLL cells from pre- and post-transplant timepoints were less intermixed within clusters (p=0.02; **Figure 2E**) and displayed a greater magnitude of gene expression change between the timepoints (p=0.02; **Figure 2F**). Similar findings were also obtained with an alternative analysis pipeline (Seurat), demonstrating that these transcriptional shifts are not dependent on the clustering algorithm (*18*) (**Figure S2A-C**). Thus, both genetic and scRNA-seq analyses indicate that early relapses comprise a heterogeneous population that is static over time, evidenced by a lack of genetic or transcriptional evolution during relapse after allo-HSCT.

The distinct post-transplant clusters seen in the late relapse patients also suggest the presence of multiple pathways of acquired resistance (i.e. cluster 6 in patient 5328 and clusters 4 and 5 in patient 5341). To delineate these pathways, we used pathway and gene set overdispersion analysis (PAGODA) to test annotated gene sets for coordinated variability across all CLL cells (*19*). Significantly overdispersed gene sets with similar cell-separation patterns were then combined to form a single ‘aspect’ of transcriptional heterogeneity. PAGODA revealed 5 major aspects of heterogeneity that distinguished the 6 CLL clusters, corresponding to ribosomal biogenesis, antigen presentation, apoptosis regulation, proliferation, and calcium/cAMP signaling (**Figure 3A**). In this manner, post-transplant pathways unique to late relapses could be identified. For example, in patient 5328, relapse was best characterized by overdispersion of apoptosis regulation, manifesting as downregulation of pro-apoptotic genes (e.g. *TP53, DFFA, BAX*) and upregulation of anti-apoptotic and cytoprotective genes (e.g. *MTRNRL2, MTRNRL8*) (*20, 21*). We also found loss of *BACH2*, a known tumor suppressor in B cell malignancies (*22*) in relapsed cells along with gain of known B cell leukemia-related pro-survival and anti-apoptotic genes (*PIM2(23), MCL1* [pre vs. post, p<2.2 × 10^−16^]) (**Figure 3A, B; S3A**). In contrast, relapse in patient 5341 was not only associated with a distinct overdispersed gene set (cluster 4; calcium/cAMP signaling evidenced by *TXNIP, DUSP1, ADD1*, and *FOS*) but also displayed intra-leukemic heterogeneity between clusters 4 and 5, especially regarding apoptosis-related genes, suggesting inter- and even intra-leukemic diversity of pathways leading to late relapse (**Figure 3B**).

**Fig. 3.**
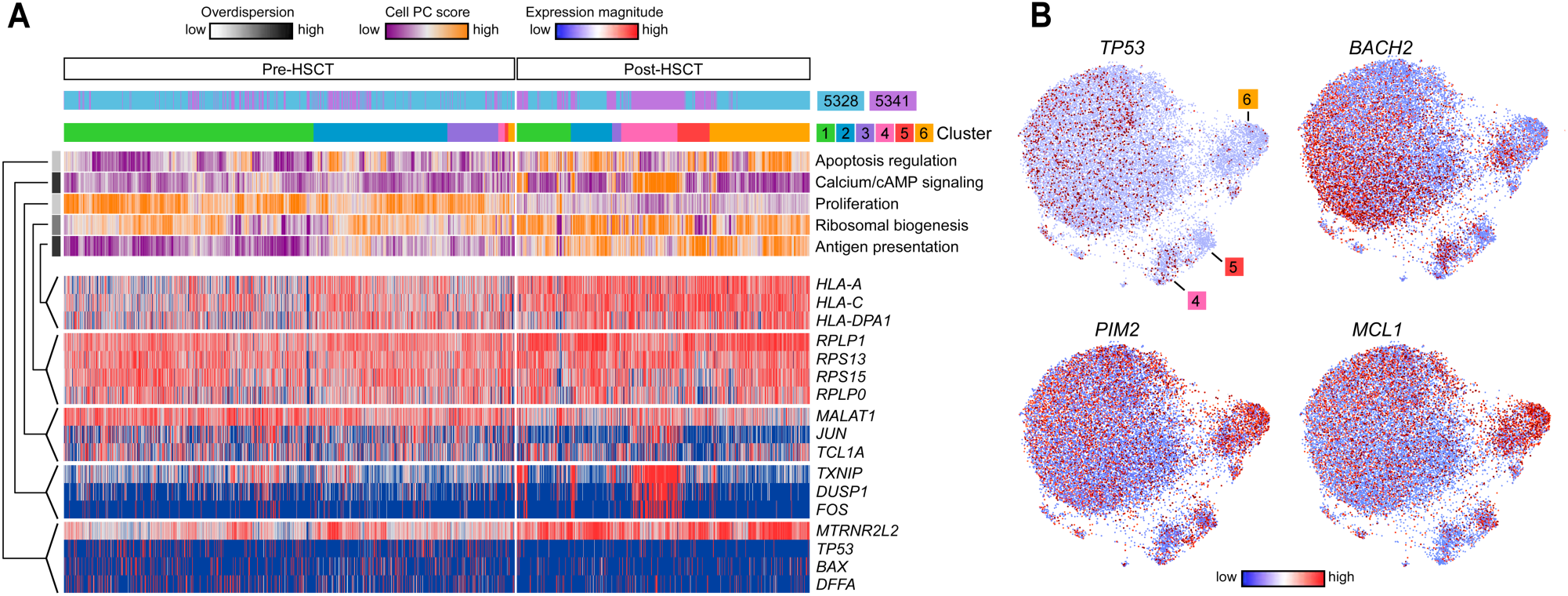
Transcriptional programs define inter- and intra-leukemic heterogeneity during late relapse. (**A**) Cells (columns) from patients 5328 and 5341 are organized by cluster assignment. For each cell, relapse kinetics, timing and principal component (PC)/aspect score are displayed. For each aspect, overdispersion score is shown by white/black color bar. Row labels summarize key functional annotations of gene sets for each aspect. For each aspect, gene expression patterns of top-loading genes are shown. (**B**) Joint graphs of CLL cells are visualized to demonstrate differential downregulation of tumor suppressor genes (e.g. *TP53* and *BACH2*) and differential upregulation of oncogenic signalling (e.g. *PIM2, MCL1*) after allo-HSCT among late relapse clusters 4, 5 and 6.

### HLA dysregulation provides no selective advantage for post-HSCT relapse of CLL

We also identified shared pathways among post-transplant populations in both late relapses (i.e. clusters 4, 5 and 6). Specifically, an aspect defined by genes involved in antigen presentation was shared among all three post-transplant, late relapse clusters (**Figure 3A, 4A**). Because loss of HLA expression has been found to contribute to relapse of myeloid leukemia after transplant (*5, 6*), we investigated further the dynamics of HLA expression variability during relapse. From our single cell data, we unexpectedly detected expression of HLA class I and II genes on post-transplant CLL cells despite relapse (**Figure 4A-B**), Moreover, we observed a wide range of HLA expression by CLL cells at both pre- and post-transplant timepoints in the late relapses. Analysis of the probability distribution of HLA class I or II genes demonstrated *increased* HLA expression for these two patients (**Figure 4C, left**, p<10^−14^ for all comparisons). Indeed, we could see that the lowest 2.5% of HLA-expressing CLL cells failed to expand during relapse in both late relapse patients 5328 and 5341, with these HLA ‘low’ cells comprising mostly pre-transplant cells (**Figure 4C, right**). At the bulk level, except for one sample with loss of HLA-A, we did not detect any somatic alterations in HLA class I, *B2M* or other genes involved in the antigen presentation pathway. Moreover, while baseline pre-HSCT bulk HLA class I gene expression in CLL was lower than observed in transplant-naïve samples, baseline bulk HLA class II expression was similar (**Figure 4D**). Notably, transcriptional downregulation of HLA class I or II genes during relapse was not observed. These data suggest that further HLA dysregulation provides little selective advantage during CLL relapse after transplant, even when present in pre-transplant subpopulations. Thus, unlike relapses from myeloid disease after transplant, genetic and/or transcriptional alterations in HLA genes are not common in the setting of CLL relapse.

**Fig. 4.**
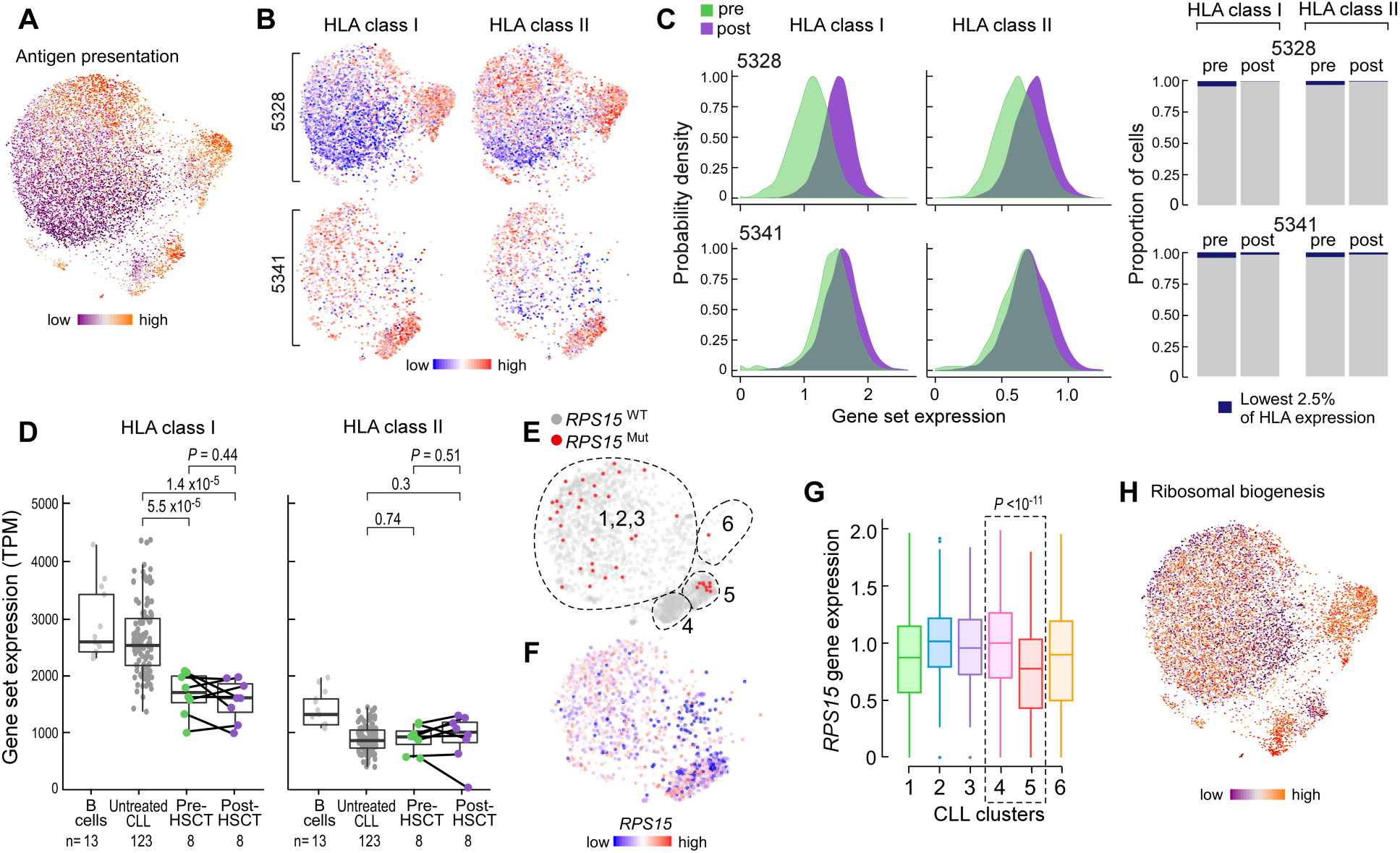
scRNA-seq analysis of late relapse clusters highlights unique features of post-transplant CLL cells. (**A**) CLL joint graph colored by individual cell score for the aspect defined as antigen presentation. (**B**) CLL joint graph displaying only cells from the indicated late relapse patient and their associated expression of HLA class I or II genes. (**C**) Left, probability densities of the range of HLA class I or II gene expression values. *P* <10^−14^ for all four comparisons, by two-sample Kolmogorov-Smirnov test. Right, stacked barplots indicating the lack of expansion of HLA class I or II ‘low’ expressing cells after allo-HSCT. *P* values for contraction determined from Fisher’s exact test. (**D**) Bulk gene set expression values for HLA class I (left) and II (right) genes from purified normal B cells, untreated CLL, and paired pre- and post-HSCT CLL cells. *P* value determined from Student’s t test (paired t test for pre- vs post-HSCT). (**E, F**) Joint graph of CLL cells from patient 5341 colored by *RPS15* mutation status (**E**) or gene expression (**F**). Dotted lines indicated approximate cluster boundaries. (**G**) Box plots showing *RPS15* gene expression by cluster for cells from patient 5341. Cluster 4 vs 5 (dotted box), *P* value calculated from two-sided Wilcoxon rank sum test. (**H**) Joint graph of all CLL cells colored by the aspect defined as ribosomal biogenesis.

### Alterations in genotype can define transcriptional changes in late relapses

The marked transcriptional change at the single cell level demonstrated by late, rather than early, relapses prompted the question of whether these transcriptional phenotypes may be driven by genetic changes. We previously reported a lack of concordance between transcriptional and genetic changes in transplant-naïve CLL cells, suggesting the presence of convergent evolution (*24*). For patient 5341, we could identify the presence of a putative CLL driver mutation in the ribosomal gene *RPS15* (p.G139C) through WES analysis; we were also able to detect this mutation within the corresponding samples’ scRNAseq data. Consistent with the bulk WES analysis, we detected a subset of cells expressing this *RPS15* mutation at the pre-transplant timepoint (clusters 1, 2, and 3) (**Figure 4E**). However, within post-transplant cells, we identified two distinct expression states corresponding to absence (cluster 4) or presence (cluster 5) of the mutation (**Figure S3C**, *P*<0.01), with associated differential expression of the calcium/cAMP signaling aspect (**Figure S3D**) as well as *TP53* and *BACH2* tumor suppressors (*P*<0.01, cluster 4 versus 5).

Upon closer examination of the post-transplant clusters, we observed higher relative expression of the *RPS15* gene (**Figure 4F, G**), along with evidence of increased ribosomal activity (**Figure 4H**), within cluster 4 relative to cluster 5. These data link the presence of the mutation with lower *RPS15* gene expression and diminished ribosomal biogenesis; conversely, absence of the mutation is associated with higher RPS15 expression and increased ribosomal activity. Together, these data demonstrate that genetic variants can segregate distinct expression states in the post-HSCT relapsed setting unlike in transplant-naïve samples, and that late CLL relapse may undergo selection pressures favoring divergent, not convergent, evolutionary paths.

### Methylome instability is unique to late relapses after allo-HSCT

We previously demonstrated locally disordered methylation (measured by the percentage of discordant reads (PDR)) as an epigenetic mechanism of genetic variability within CLL, with increased PDR associated with a more aggressive clinical course in transplant-naïve CLL (*25*). We therefore investigated whether epigenetic dysregulation could underlie the observed phenotypic changes by comparing the change in PDR over time between early and late relapses (**Table S10**). The change in PDR for early relapses was minimal, while late relapses exhibited greater changes in PDR across multiple genomic elements (**Figure 5A**). By contrast, PDR did not change within CLL samples relapsing after chemotherapy alone, matched for timing with late relapses after transplant (**Figure S3C, 5A**). When we modeled the quantitative changes in PDR as a function of time, we confirmed a moderate increase in the rate of change of PDR for the late relapsers compared to the chemotherapy-treated subjects, suggesting time between samples was not a contributing factor to our PDR measurements (**Figure 5B**, Bayes factor=1.5) (*26*). Altogether, these findings are consistent with the notion of post-HSCT immune pressure selecting for increased intra-leukemic methylation variability as an underlying mechanism of phenotypic changes. Indeed, genes with increased promoter PDR in late relapses were enriched for multiple stem cell pathways, implicating a common stem-like state that is acquired during late relapses through increased methylome instability (**Figure 5C-F, Table S11**) and is consistent with the transcriptional upregulation of similar modules in this group (**Figure 1E**).

**Fig. 5.**
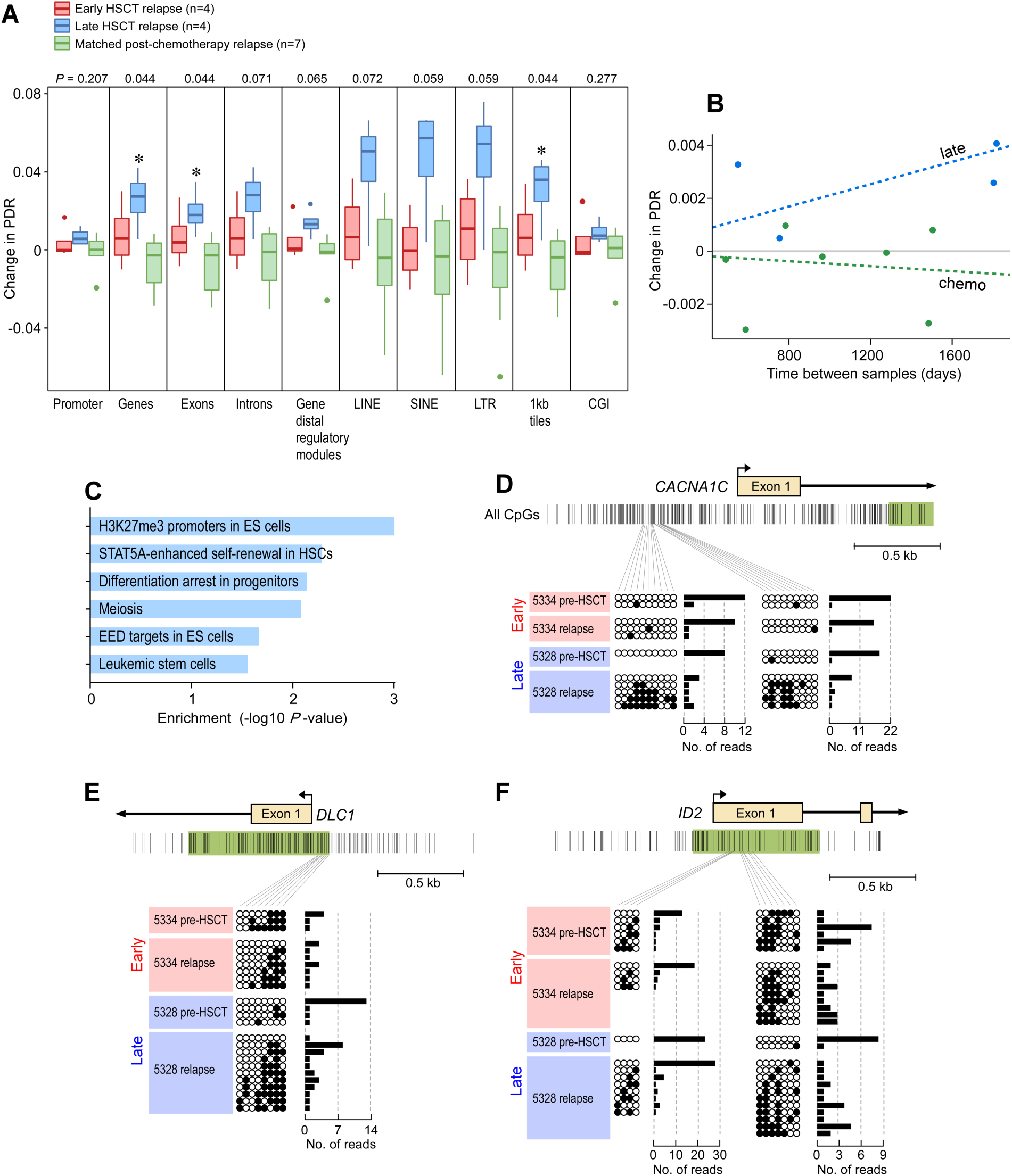
Methylome instability characterizes late relapse after allo-HSCT. (**A**) Boxplots of the change in locally disordered methylation as measured by the percentage of discordant methylated reads (PDR). Shown are values for early relapse (n=4), late relapse (n=4) and matched post-chemotherapy treated relapse patients (n=7). * = p<0.05 determined from Kruskal Wallis test. (**B**) Graph depicts change in PDR as function of change in time to progression. Dotted line represents Bayesian linear model fitted to each group. Points represent weighted averages of all genomic regions per patient. Bayes factor=1.5. (**C**) Stem cell gene sets are enriched from sites of significant methylation differences (absolute change >10%) between pre-HSCT and relapse samples in late relapse patients. *P* values were calculated from Fisher’s exact test. (**D-F**) Ordered and disordered methylated reads for patients 5334 (early relapse) and 5328 (late relapse), respectively, for three stem cell gene-associated promoters.

## Discussion

We have investigated leukemic cell-intrinsic factors that contribute to GvL resistance by studying CLL relapse after transplant. We find that underlying genetic and epigenetic trajectories shape the kinetics of leukemic relapse. Indeed, early relapses after allo-HSCT were marked by a molecular stability that was remarkably consistent across genetic, transcriptional and epigenetic measurements. In contrast, late relapses exhibited striking genetic and epigenetic change along with evidence of neoantigen depletion, consistent with manifest single cell transcriptional shifts that were unique to each patient.

In addition, these evolutionary paths appeared unique to the transplant setting. For example, unlike in transplant-naïve CLL samples, we found a post-HSCT relapse to exhibit concordant genotype-phenotype relationships that defined transcriptional changes and suggested divergent evolutionary paths. Furthermore, in contrast to post-chemotherapy relapses exhibiting markedly stable methylomes, recurrences late after allo-HSCT were characterized by widespread methylome instability across the epigenome. These data suggest that the evolutionary consequences of the GvL bottleneck may in fact be distinct from those imposed by chemotherapeutic ones and that therapeutic strategies to prevent or treat cancer recurrence after such immunologic bottlenecks should account for this potential diversification.

Our transcriptional and epigenetic data implicate stem cell pathways as contributory for CLL relapse after allo-HSCT given their enrichment in early relapsed CLL cells pre-HSCT and their acquisition in late relapsed CLL cells post-HSCT. Indeed, immune evasion and cell mobility are well-studied properties of stem cells (*27–29*). While the upregulation of pathways such as BCR and FcR signaling during late relapse suggest additional drivers of GvL resistance, our results highlight stem cell properties for future investigation as putative predictors of relapse and a mechanistic basis for GvL escape.

These studies demonstrate the applicability of single cell genomics to delineate genetic selection pressures for leukemic subpopulations within individual patients. For example, in contrast to previous reports in myeloid malignancies (*5, 6*), we could see no role for HLA gene downregulation during leukemic evolution. In fact, scRNA-seq enabled a high-resolution portrait of the static and even contracting dynamics of such alterations, demonstrating that even when they are present in pre-transplant subsets, no subsequent evolutionary advantage exists. These data suggest that the rules of GvL resistance differ depending on the leukemic subtype.

Although our study focused on leukemic cells, understanding how immune cells, particularly T cells (*30*), also co-evolve during post-HSCT relapse is critical for reversing GvL resistance. Moreover, future studies should examine how the site of disease and the spatial microenvironment can impact these co-evolutionary interactions given their known influence on CLL behavior (*31*). Finally, although our study focused on CLL, these results should motivate similar efforts within other lymphoid lineages where additional factors modulating GvL resistance may be discovered. Nevertheless, our results in studying CLL in this context highlight the roles of stem cell genes and evolutionary dynamics in promoting relapse as well as suggest biological pathways for future investigation as putative predictors of GvL escape.

## Materials and Methods

### Study Design

The overall objective of this study was to identify leukemia-intrinsic features that shape CLL relapse after allo-HSCT. The cohort size was determined by sample availability, and we identified 10 patients treated for CLL at the DFCI that had paired, pre- and post-allo-HSCT samples available. Patients were consented to an institutional review board (IRB)-approved research protocol (01-206). The specimens were collected after informed consent under a study protocol that was approved by the DFCI Human Subjects Protection Committee and conducted in accordance with the Declaration of Helsinki. We undertook a comprehensive assessment of genetic, epigenetic and single cell transcriptomic features from purified CLL cells and investigated longitudinal changes in a pair-wise fashion. All samples and data were included in our analyses. Primary data is deposited in dbGaP, and results of primary analyses are included in **Tables S1-S11**.

### Patients and samples

As described above, a total of twenty paired CLL samples were obtained pre- and post-HSCT from patients treated at the Dana-Farber Cancer Institute (DFCI) between 2006-2016. 18 of 20 samples came from cryopreserved peripheral blood mononuclear cells (PBMCs), 1 from cryopreserved bone marrow mononuclear cells (BMMCs), and 1 from a formalin-fixed (FFPE) bone marrow biopsy (see **Table S1**). Seven paired CLL blood samples were obtained pre- and post-chemotherapy from patients enrolled on clinical research protocols approved by the Human Subjects Protection Committee at UCSD (CLL Research Consortium). PBMCs and BMMCs from all patients were isolated by Ficoll-Hypaque density gradient centrifugation, cryopreserved with 10% dimethyl sulfoxide, and stored in vapor-phase liquid nitrogen until time of use. For all samples with at least 2×10^6^ suspension cells (n=18 of 20 DFCI samples; n=14 of 14 UCSD samples), tumor cells were purified by flow cytometric isolation of CD19^+^/CD5^+^ cells for downstream nucleic acid extraction.

### *IGHV* and prognostic marker analysis

As per convention, *IGHV* unmutated status was defined as greater than or equal to 98% homology to the closest germline match and *IGHV* mutated if homology was below this cut-off. Assessments for the cohort were done at the CLL Research Consortium Tissue Core (UCSD, San Diego) or at Integrated Oncology (New York, NY) as described in detail elsewhere. When available, routine pathological assessment of common CLL chromosomal abnormalities was included based on interphase FISH with Vysis probes (Abbott Molecular, Des Plaines, IL).

### Whole-exome sequencing data generation and preprocessing

For whole-exome sequencing (WES), we used standard Broad Institute protocols as previously described (*12*). DNA of all samples was isolated using the All-prep DNA/RNA Mini kit (Qiagen Hilden, Germany). Tumor DNA was derived from CLL cells, and matched germline line DNA came from both pre-transplant CD4^+^ T cells as well as donor DNA derived from the transplant product. Tumour and normal DNA concentration were measured using PicoGreen® dsDNA Quantitation Reagent (Invitrogen, Carlsbad, CA). A minimum DNA concentration of 5 ng/µL was required for sequencing. All Illumina sequencing libraries were created with the native DNA. The identities of all tumour and normal DNA samples were confirmed by mass spectrometric fingerprint genotyping of 95 common SNPs by Fluidigm Genotyping (Fluidigm, San Francisco, CA). Sequencing libraries were constructed with the Illumina Rapid Capture Enrichment kit. Pooled libraries were normalized to 2nM and denatured using 0.2 N NaOH prior to sequencing. Flowcell cluster amplification and sequencing were performed according to the manufacturer’s protocols using either the HiSeq 2000 v3 or HiSeq 2500. Each run was a 76 bp paired-end with a dual eight-base index barcode read. Standard quality control metrics, including error rates, percentage-passing filter reads, and total Gb produced, were used to characterize process performance before downstream analysis. Data analysis included de-multiplexing and data aggregation.

#### Mutation calling

We used the Getz Lab CGA WES Characterization pipeline [https://docs.google.com/document/d/1VO2kX_fgfUd0X3mBS9NjLUWGZu794WbTepBel3cBg08/edit] developed at the Broad Institute to call, filter and annotate point mutations, insertions and deletions. Paired-end Illumina reads were aligned to the hg19 human genome reference using the Picard pipeline [https://software.broadinstitute.org/gatk/documentation/tooldocs/4.0.1.0/picard_fingerprint_CrosscheckFingerprints.php https://software.broadinstitute.org/gatk/documentation/tooldocs/4.0.0.0/picard_analysis_CollectMultipleMetrics.php]. Cross-sample contamination was assessed with the ContEst (*32*) tool (5% threshold). Point mutations and indels were identified using MuTect(*33*) and Strelka(*34*), followed by annotation using Oncotator (*35*). Possible artifacts due to orientation bias, germline variants, sequencing and poor mapping were filtered using a variety of tools including Orientation Bias Filter (*36*), MAFPoNFilter (*37*), and RealignmentFilter. All somatic alterations thus identified were further manually inspected in IGV (*38*) and apparent false positives were removed.

#### Somatic copy number alteration identification

Copy number events were called and filtered using GATK4 ModelSegments (*39*) [https://gatkforums.broadinstitute.org/dsde/discussion/11682/; https://gatkforums.broadinstitute.org/dsde/discussion/11683]. In order to minimize false positives, we utilized a copy number panel-of-normals created based on germline samples processed using the same platform. We applied a custom conversion script to format the outputs of ModelSegments (both copy ratio and allelic fraction) to be compatible with ABSOLUTE (*39, 40*), the tool used to estimate sample purity and ploidy as well as cancer cell fractions (CCFs). ABSOLUTE solutions were picked by manual inspection. The final chosen purity and ploidy solutions were used to estimate CCFs for detected somatic alterations in each sample.

#### Evaluation of clonal evolution

Inference of clonal structure, phylogenetic relationship between clones and evolution between pre- and post-treatment time points within a sample was performed using the PhylogicNDT tool (*11*). For each PhylogicNDT assigned cluster within an individual, pairs of CCF values were drawn at the two points using their respective marginal posterior pdfs (N=10,000). The posterior probability that a cluster was evolving was estimated as the proportion of draws where the difference in CCF values was < 0.2. The Benjamini-Hochberg FDR correction method was used to account for multiple hypothesis testing for all clusters within an individual. An individual was considered to be evolving if there was at least one cluster with adjusted P value <0.1. Clusters were defined as ‘contracting’ or ‘expanding’ depending on the corresponding sign of the change in CCF.

#### HLA and neoantigen prediction

We used Polysolver (*41*) for computational allele inference of major MHC class I genes from normal exome sequencing data (HLA-A, HLA-B and HLA-C). We predicted binding affinities for all possible 9 and 10-mers arising from coding mutations, insertions or deletions against all HLA alleles for a given patient using NetMHCpan-4.02 (*42*). Strong and weak affinity neoantigen binders were identified based on binding scores (≤ 50 nM for strong, ≤ 500 nM for weak), and those with undetectable gene expression levels were removed.

### RNA sequencing data generation and analysis

A cDNA library was prepared from poly-A selected RNA and sequenced on an Illumina platform. Paired-end transcriptome reads were mapped to the UCSC hg19 reference genome using STAR. Differential expression analyses were conducted using *DESeq2* R package v.1.26.0. Differentially expressed genes were identified using a cutoff of FDR p ≤0.25 and subsequently used for gene set enrichment analysis (GSEA) of the C2 subset of the Molecular Signature Database. Differentially enriched gene sets were identified using a cutoff of FDR p ≤0.25.

### Single cell transcriptome generation and analysis

#### Sample Preparation and Processing

Cryopreserved samples were thawed the day of experiments. Dead cells were removed via an Optiprep selection protocol such that cells collected just below the PBS layer were >95% viable on trypan blue staining (*16*). Viable cells were subjected to immunomagnetic selection (MACS CD19 MicroBeads; Miltenyi Biotec, cat. No. 130-050-301), using MS columns, to isolate CD19+ and CD19-cells. Cells were prepared at a concentration of 1.15 × 10^5^ cells/mL in a 15% OptiPrep solution in PBS and kept on ice until time of encapsulation.

Cell encapsulation was performed in a light-protected environment using the Olympus CKX53 microscope, Point Grey Chameleon3 1.3 MP Mono USB3 Vision camera, and OEM Syringe Pump Modules. RT/lysis mix and barcoded hydrogel beads (BHBs; from 1CellBio (Watertown, MA)) were prepared as previously described (*16*) and directly before time of encapsulation. During encapsulation, cells and RT/lysis mix were kept at 4°C in their respective syringes using refrigerated copper coiling, and the collection tube was kept cool on an ice rack. All reagents were simultaneously loaded onto microfluidic devices (microfluidic chip; 1CellBio (Watertown, MA)). The Harvard Apparatus Pump Software and the Point Grey camera software were used to control the parameters of encapsulation with the following working flow rates: 50-70 µL/hr for BHBs, 340-380 µL/hr for carrier oil, 250 µL/hr for cell suspension, and 250 µL/hr for RT/lysis mix, which produced a bead occupancy of 70-85% and droplets sizes ranging from 2.5 to 3.5 nL. Encapsulation time for 3,000 cells was approximately 9 to 12 minutes with a calculated cell doublet percentage of 5.45% to 7.22%.

#### Library Preparation and Sequencing

Library preparation and sequencing proceeded as previously published (*16*). Briefly, after the target number of cells was encapsulated, the emulsion of droplets was exposed to UV light (365 nm at ∼10 mW/cm2, UVP B-100 lamp) for 8 minutes on ice. The emulsion was incubated at 50°C for 2 hr, 70°C for 15 min, and 4°C for 1 min before splitting into 3,000 cell aliquots. The emulsion was broken using 10 µL of 20% (v/v) 1H,1H,2H,2H-Perfluorooctanolin in Novec-7500 and stored in -80°C until libraries were ready to be processed. In vitro transcription was performed overnight for 16 hours at 37°C. Fragmentation of amplified RNA libraries was performed for 1.5 minutes at 70°C. The final sequence-ready libraries were amplified by 7-9 PCR cycles on average as determined by a diagnostic qPCR. Libraries were quantified by Agilent BioAnalyzer and multiplexed with unique PCR indices in sequencing batches of up to 12,000 cells. The NextSeq Illumina Sequencer was used to sequence libraries using custom sequencing primers.

#### scRNA-Sequencing Data Analysis

Raw count data were analyzed using the dropEst software suite(*43*). The dropTag tool was used to annotate reads with cellular barcodes and collapse the different read files. Reads were aligned to the hg38 genome using tophat2 (*44*) provided with gene annotations from UCSC. Read quantification was performed using dropEst using the simple correction model (-m) and correctional correction of UMI errors (-u).

The panel of samples was jointly analysed using the Conos suite(*17*) and pagoda2 packages (https://github.com/hms-dbmi/pagoda2). Briefly, normalization of individual datasets was performed using pagoda2, admitting cells with at least 500 molecules. Conos integration was performed using PCA rotation space, calculating top 30 PCs on a union of 2000 overdispersed genes for each pair of datasets. LargeVis embedding was used to visualize the joint subpopulations. Tumor cells were manually annotated as the CD19+/CD5+ cluster. The tumor cells were then re-integrated separately using Conos using CPCA rotation with 30 PCs. The resulting joint graph of CLL cells was visualized using UMAP embedding. Clusters within the CLL cells were then determined using the Leiden algorithm, with the resolution of r=0.7. Higher resolution clustering (r=1.9) was then selected to annotate distinct subpopulations appearing in late relapse samples (clusters 4-6).

An unsupervised clustering analysis was also performed using the *Seurat* R package v3.1.1(*18*). In particular, an integrated approach was taken in order to minimize individual patient effect yet maintain differences in timing. Principal component analysis (PCA) dimensionality reduction was performed with the highly variable genes as inputs. The PCs were then used to calculate a uniform manifold approximation and projection (UMAP) for the integrated data. Clusters were called at a resolution of 1 using the first 30 principal components. Tumor cell clusters were manually selected via CD5+/CD19+ and integrated again without input from additional immune cells from patients. Clusters on this tumor-specific data were called at a resolution of 0.5 using the first 30 principal components.

### Methylome sequencing and analysis

RRBS libraries were generated from 25-100 ng of input DNA using the Ovation Methyl-Seq System (NuGen) following the manufacturer’s recommendation. These reads were aligned to the human hg19 genome using BSmap with flags -v 0.05 -s 16 -w 100 -S 1 -p 8 –u. An average of 23.1M reads per sample were aligned correctly. An average of 5.4M CpGs were covered per sample. The methylation state of each CpG was determined by comparing bisulfite-treated reads aligning to that CpG with the genomic reference sequence. The methylation level was computed by dividing the number of observed methylated cytosines (which did not undergo bisulfite conversion) by the total number of reads aligned to that CpG. The proportion of discordant reads (PDR) was calculated for each CpG as previously described(*25*). Gene set enrichment analysis was limited to the C2 MSigDB gene set collection, available at: https://www.gsea-msigdb.org/gsea/msigdb/genesets.jsp?collection=C2. Gene set enrichment analysis was performed for genes that exhibit consistently elevated PDR (greater than mean promoter PDR of 0.1 in >50% of samples), and a Fisher’s exact test was used to measure the enrichment of these gene sets in late relapses (compared to early relapses). Statistical analysis for methylome data was performed with R version 3.4.0.

### Statistical analysis

Statistical analysis was performed with R version 3.5.3, unless otherwise specified. Categorical variables were compared using the Fisher Exact test, and continuous variables were compared using the Student’s t-test, Wilcoxon rank sum test, or Kruskal-Wallis test as appropriate. Probability densities were estimated using kernel density estimations, implemented by the geom_density function within the ggplot2 package, and compared using the Kolmogorov-Smirnov test. A Bayesian linear regression model was calculated via the MCMCregress function within the MCMCpack package. The Bayes factors of these were calculated using the ecdf function native to R.

#### Analysis of cell intermixing

To quantify the degree of intermixing between pre- and post-transplant cells for each patient, we calculated the proportion of neighbors belonging to the same time point as the cell of interest. The nearest neighbors of each cell were determined as 500 closest cells on the existing embedding. The mean proportions were calculated for each patient and compared using a two-sided Welch t-test. A similar calculation was done within Seurat using the FindNeighbors function and the integrated nearest neighbors matrix along with a neighborhood size of 100 cells.

#### Quantification of gene expression distance

To compare the extent of expression difference in tumor cells between pre- and post-transplant timepoints for the patients with early and late relapse, for each patient we sampled a random subset of a 100 cells from each timepoint, and then calculated the expression distance as a Jensen-Shannon divergence between the vectors for the two timepoints. The values in each vector give the probability of observing a molecule of a given gene in a given time point. The statistical significance of the mean inter-timepoint distance between early and late relapse patients was assessed using a one-sided Welch t test.

## Acknowledgments

We are grateful for members of Allon Klein’s laboratory as well as Colin Brenan, Mike Brenan, and Ilke Akartuna of 1CellBio for discussions and advice regarding setup of the inDrops platform. We also thank the study nurses and clinical staff that banked these samples, and the patients who generously consented for the research use of these samples. Finally, we appreciate members of the Wu laboratory for their feedback and support. Funding: This work was supported in part by the NCI (1R01CA155010, P01CA206978, U10CA180861). P.B. was supported by a Physician-Scientist Training Award from the Damon Runyon Cancer Research Foundation, an Amy Strelzer Manasevit Scholar Award from the Be The Match Foundation, and an American Society of Hematology Fellow Scholar Award. S.H.G. is supported by a Kay Kendall Leukaemia Fund fellowship. S.A.S is supported by a grant from the NCI (R50CA211482). D.N. is supported by a grant from the NIH (5P30 CA006516). P.V.K. is supported by a grant from the NHLBI (1R01HL131768). C.J.W. is a Scholar of the Leukemia and Lymphoma Society.

## Author contributions

P.B. conceived and supervised the study, designed and performed experiments, analyzed and interpreted results, and wrote the manuscript. C.E. analyzed genomic and transcriptomic data and edited manuscript. V.N.N. adapted the inDrops platform, performed flow cytometric sorting of CLL cells and generated scRNA-seq data. K.C. analyzed and interpreted methylome data. S.H.G. analyzed and interpreted scRNA-seq data and edited manuscript. S.A.S. analyzed and interpreted whole exome sequencing data. J.F. analyzed and interpreted whole exome sequencing data. N.B. processed and analyzed scRNA-seq data. N.B. curated patient data. S.F., L.E., I.L., and G.A.G contributed to analysis of whole exome sequencing data. L.Z.R and T.J.K performed IGHV sequencing. A.W.M. generated the bulk RRBS libraries with supervision from A.G. J.R.B. oversaw patient care and contributed samples. V.T.H. oversaw patient care, provided clinical data from the BMT repository and contributed to interpretation of clinical data. D.N. contributed to statistical analysis and edited manuscript. R.J.S. oversaw patient care, contributed samples, interpreted results and edited manuscript. J.R. oversaw patient care, contributed samples, interpreted results and edited manuscript. E.P.A. oversaw patient care, contributed samples and interpreted results. P.V.K. designed, performed, and interpreted analyses of scRNA-seq and methylome data and edited manuscript. C.W. conceived and supervised study, interpreted results and edited manuscript.

## Data and materials availability

Exome, transcriptome, single cell transcriptome, and methylome data will be submitted to NCBI’s Database of Genotypes and Phenotype (dbGaP; https://www.ncbi.nlm.nih.gov/gap) under study number: [pending].

## Supplementary Materials

**Fig. S1.**
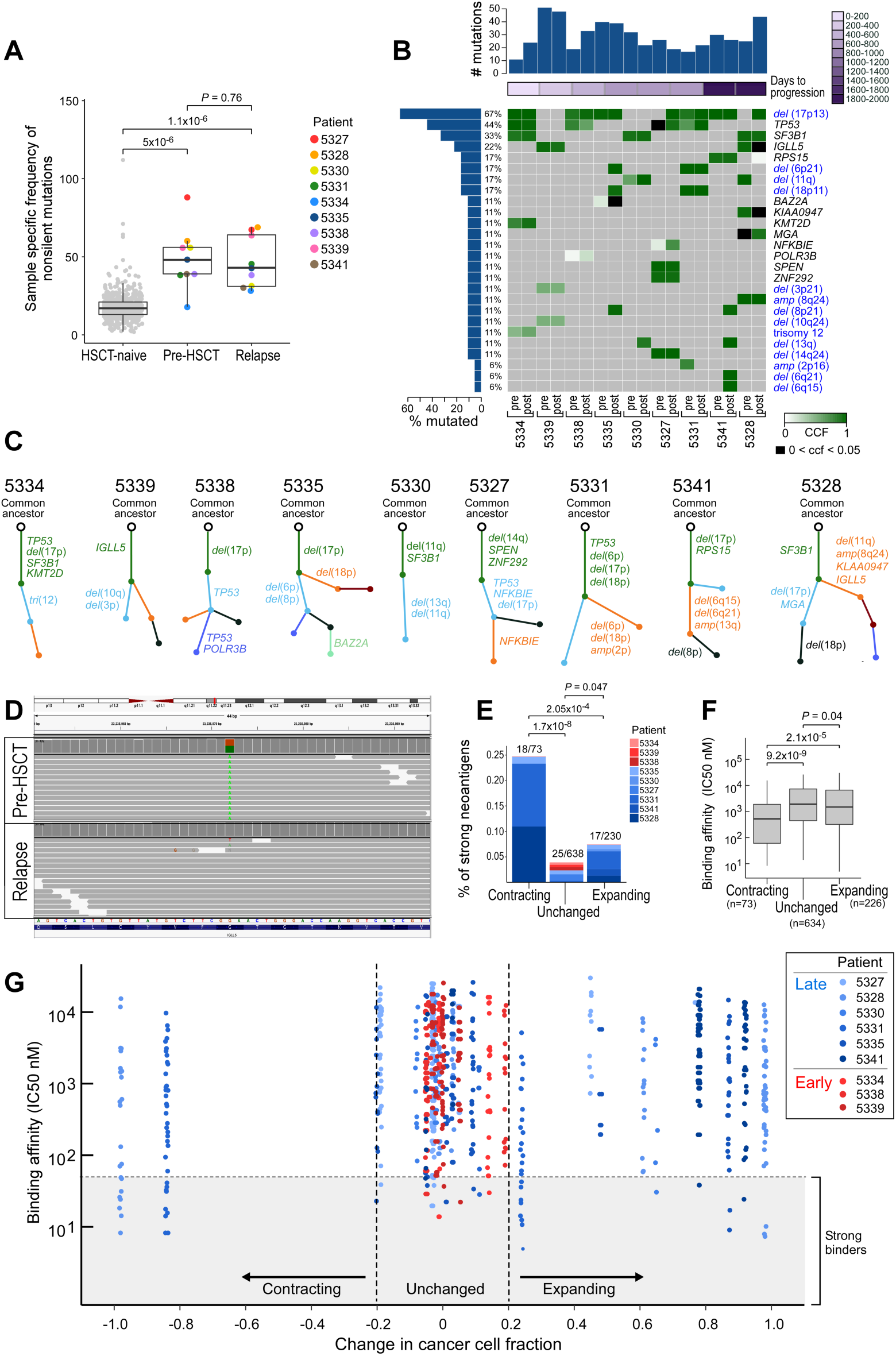
Frequencies of mutations, CLL cancer drivers and neoantigens. (**A**) Boxplot of the sample specific frequency of non-silent mutations in treatment-naïve CLL samples as compared to pre- and post-relapse. *P* values determined from Wilcoxon ranked sum test. (**B**) Comut plot representing change in cancer cell fraction of various somatic single nucleotide variants and copy number alterations (blue font) (rows) for 9 paired pre-/post-RIC patients (columns). Time to progression is shown by the purple color bar. Bar plot showing total number of mutations per patient per timepoint across top, as well as a bar plot showing percent of samples containing an individual mutation across left. Copy number alterations shown in blue. (**C**) Inferred phylogenetic structure for each patient using PhylogicNDT. Colors represent distinct mutational clusters. (**D**) Snapshot from the Integrated Genomics Viewer of a contracting neoantigen predicted from a mutation in the *IGLL5* driver gene (g.chr22:23235972G>A), detectable in the pre-HSCT track (top) but not in the relapse track (bottom). (**E**) Stacked bar plot indicating the percentage of strong neoantigens (IC50 ≤ 50nM) among contracting, unchanged, or expanding neoantigens and the proportion contributed by each patient, colored by early or late relapse timing. Total number of strong neoantigens indicated at top of bar for each neoantigen class. *P* value determined from Fisher’s exact test. (**F**) Box plot indicates median binding affinity for all predicted contracting, unchanged or expanding neoantigens. *P* value determined from one-sided Wilcoxon ranked sum test. **(G)** Scatterplot of change in cancer cell fraction versus binding affinity for all neoantigens. Color denotes individual patients, with shades of red representing early and shades of blue representing late relapse after allo-HSCT. Dotted line denotes absolute change in CCF of ≥0.2. Dashed line denotes threshold for strong neoantigens, defined by an IC50 ≤ 50nM.

**Fig. S2.**
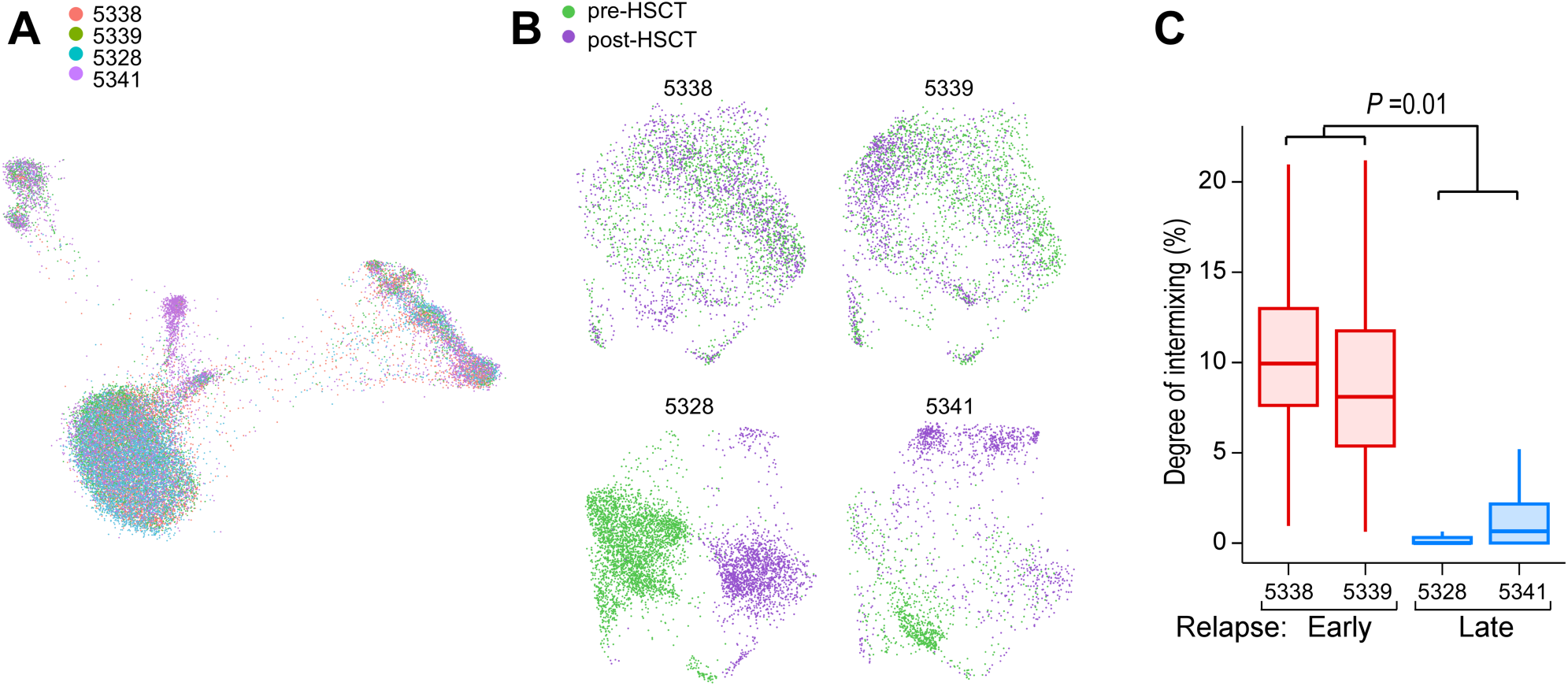
Single cell transcriptome analyses. (**A**) UMAP joint graph (with Conos) of all cells from all patients, colored by patient. (**B**) Individual UMAP clusterings (with Seurat) of all CD19^+^CD5^+^ CLL cells for each patient, colored by timing. (**C**) kNN-based quantification of timepoint intermixing across clusters, identified by Seurat, on a per-cell basis for each patient. *P* value determined from two-sided Welch t-test comparing means of individual cell values per patient, grouped into early (n=2) versus late (n=2) relapses.

**Fig. S3.**
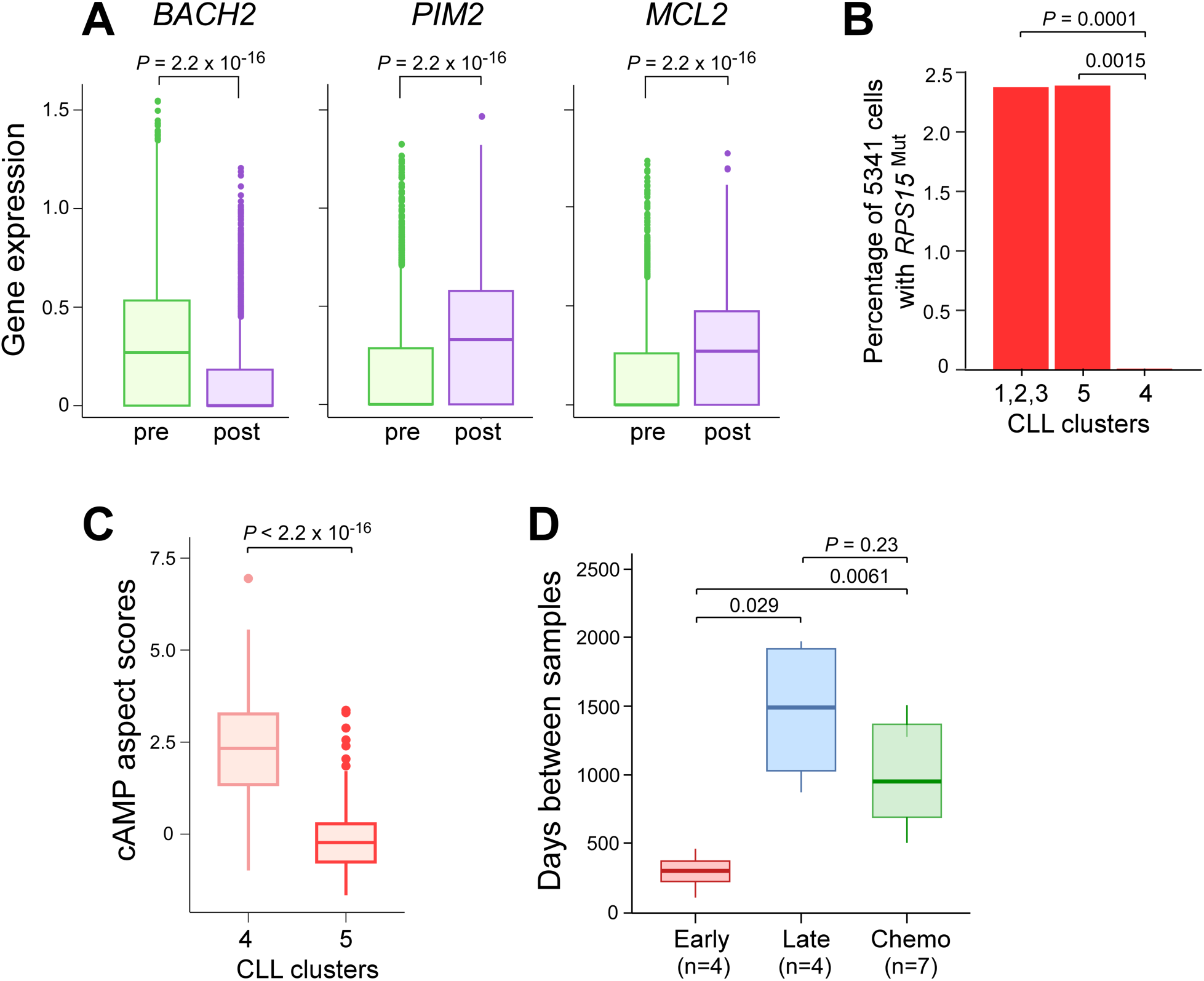
Quantification of single cell gene expression values. (**A**) Boxplots of normalized gene expression values for *BACH2, PIM2*, and *MCL1* comparing pre- to post-HSCT cells from patient 5328. (**B**) Cluster 4 contains significantly fewer RPS15^Mut^ cells than post-HSCT cluster 5 or the pre-HSCT clusters 1,2, and 3. *P* value calculated from Fisher’s exact test. (**C**) Significantly higher per-cell expression of the aspect defining cAMP signaling in cluster 4 than cluster 5. *P* value calculated from two-sided Wilcoxon ranked sum test. (**D**) Boxplot shows time between pre- and post-HSCT samples for early (n=4), late (n=4) or after chemotherapy-only relapsed groups. The *P* values were calculated from a two-sided Wilcoxon ranked sum test.

**Table S1.** Clinical characteristics.

| Patient ID | 5327 | 5328 | 5330 | 5331 | 5334 | 5335 | 5338 | 5339 | 5340 | 5341 |
| --- | --- | --- | --- | --- | --- | --- | --- | --- | --- | --- |
| Age at SCT | 41 | 55 | 56 | 47 | 53 | 59 | 54 | 63 | 70 | 59 |
| Sex | M | F | F | F | M | F | M | M | F | M |
| *IGHV* status | M | U | U | U | N/A | U | N/A | M | M | N/A |
| Disease status prior to allo-HSCT | PR | PR | PR | PR | PR | PR | PR | PR | PR | PR |
| Donor source | PB | PB | PB | PB | PB | PB | PB | PB | PB | PB |
| Donor sex | F | M | F | M | F | M | F | M | M | F |
| Donor type (matched) | Rel | Rel | Rel | Rel | Rel | Rel | Rel | Rel | Un | Un |
| Conditioning intensity | RIC | RIC | RIC | RIC | MA | RIC | RIC | RIC | RIC | RIC |
| Conditioning regimen | flu/bu | flu/bu | flu/bu/r | flu/bu/r | cy/tbi | flu/bu | flu/bu | flu/bu | flu/bu | flu/bu |
| GvHD ppx | t/m | t/m | t/m/rap/r | t/m/rap/r | t/m | t/m/rap | t/m/rap | t/m/rap | t/m | t/m/rap |
| Days from dx to SCT | 2409 | 3395 | 2884 | 1278 | 2482 | 657 | 4599 | 5950 | 4380 | 2628 |
| Days from sample to SCT | 300 | 154 | 273 | 205 | 10 | 214 | 14 | 6 | 71 | 90 |
| Days from SCT to relapse sample | 782 | 1825 | 731 | 1068 | 83 | 643 | 442 | 304 | 194 | 1801 |
| Best response to SCT | CR | CR | CR | CR | P | MR | PR | PR | CR | CR |
| Paired samples for WES | Y | Y | Y | Y | Y | Y | Y | Y | N | Y |
| Paired samples for RNA-seq | Y | Y | Y | N | Y | Y | Y | Y | N | Y |
| Paired samples for RRBS | Y | Y | N | N | Y | Y | Y | Y | Y | Y |
| Paired samples for scRNA-seq | N | Y | N | N | N | N | Y | Y | N | Y |

M=mutated, U=unmutated; PR=partial response; PB=peripheral blood; Rel=related, Un=unrelated; RIC=reduced intensity,
MA=myeloablative; flu=fludarabine, bu=busulfan, r=rituximab, cy=cyclophosphamide, TBI=total body irradiation;
t=tacrolimus, m=methotrexate, rap=rapamycin; CR=complete response, P=progression, MR=mixed response; Y=yes, N=no

Included in **Auxiliary Supplementary Materials**

Table S2. Whole exome sequencing metrics.

Table S3. Somatic single nucleotide variants and insertions/deletions

Table S4. Somatic copy number alterations (sCNAs) and ABSOLUTE-calculated cancer cell fraction estimates.

Table S5. Clustering- and phylogeny-adjusted subclone-CCFs and driver annotations.

Table S6. Neoantigens predicted per patient

Table S7. Differentially expressed genes in pre-transplant CLL cells (early versus late)

Table S8. Differentially expressed genes in late relapsers (pre-versus post-transplant)

Table S9. Single cell RNA sequencing metrics

Table S10. Promoter methylation values

Table S11. Promoter PDR values

